# Antimicrobial Potential and Characterization of Silver Nanoparticles Synthesized from *Ocimum Sanctum* Extract

**DOI:** 10.1101/2024.09.19.613829

**Authors:** Mustapha Sulaiman, Ma’aruf Abdulmumin Muhammad, Aminu Shehu Sulaiman, Amina Lawan Abubakar, Ritu Sharma, Abubakar Musa Shuaibu, Mamudu Aliyu, Ibrahim Tasi’u Mustapha, Raksha Tiwari

## Abstract

The growing resistance of microbes to conventional antibiotics and the environmental impact of chemical synthesis methods highlight the urgent need for effective and sustainable antimicrobial agents. In this study, we address these challenges by synthesizing silver nanoparticles (AgNPs) using *Ocimum sanctum* leaf extract, a green and efficient method. We confirmed AgNPs synthesis through UV-vis spectroscopy, observing a Surface Plasmon Resonance (SPR) peak centered at 420 nm. X-ray Diffraction (XRD) analysis showed intense peaks at 37.81°, 45.01°, 63.91°, and 78.0°, corresponding to Bragg’s reflections at 111, 200, 220, and 311, respectively, with an average particle size of approximately 18 nm calculated using Scherrer’s formula. The reduction of Ag^+^ was monitored using Atomic Absorption Spectroscopy (AAS), which revealed a rapid decrease from 2.73 ppm to 0.023 ppm within four minutes, driven by the active reducing agents in the *O. sanctum* extract. The Scanning Electron Microscope (SEM) image confirmed the spherical shape of the AgNPs, with an average size of 23.82 ± 4.17 nm. Fourier Transform Infrared (FTIR) spectroscopy identified absorption peaks at 1635 cm^-1^ and 3430 cm^-1^, associated with the amide I bond of proteins and OH stretching in alcohols and phenolic compounds, respectively. The synthesized AgNPs exhibited significant antimicrobial activity, demonstrated by a dose-dependent inhibitory effect against bacterial strains. The inhibition zone measured 6 mm at the lowest concentration of 5 µg/L and increased progressively to 14 mm at the highest concentration of 25 µg/L, indicating strong antimicrobial potential even at low dosages.

## Introduction

The production and utilization of silver nanoparticles (AgNPs) have attracted noteworthy interest because of their distinct physicochemical characteristics and wide range of uses in disciplines like environmental research [1], electronics [2], and medicine [3]. Among these, AgNPs are especially useful in fighting microbial infections due to their strong antibacterial qualities, which is becoming more and more important in light of the rise in antibiotic resistance [4]. The development of new antimicrobial medicines is imperative due to the serious threat that antibiotic-resistant bacteria pose to public health [5]. Because of their broad-spectrum antibacterial activity, low toxicity to human cells, and ability to get around bacterial resistance mechanisms, silver nanoparticles have emerged as a possible option [6].

Concerns regarding the sustainability and environmental effect of traditional methods for producing silver nanoparticles have been raised since they frequently entail substantial energy usage and dangerous chemicals [7]. Hazardous byproducts that pose a threat to human health and the environment can arise from the employment of toxic stabilizers and reducing agents in chemical synthesis processes [8]. In order to create nanoparticles in a more economical and environmentally responsible way, researchers have resorted to green synthesis techniques in response to these difficulties, using natural plant extracts as reducing agents [9]. In addition to lessening the impact on the environment, this biogenic manufacturing technique offers the possibility of functionalizing nanoparticles with beneficial compounds found in plant extracts [10]. The synthesis of nanoparticles using plant extracts is in line with green chemistry principles, which encourage the use of renewable resources and lessen dependency on dangerous materials [11 & 12].

*Ocimum sanctum*, a plant also referred to as holy basil or tulsi, is well-known for its therapeutic qualities, which include anti-inflammatory [13, 14], and antioxidant capabilities [15]. *Ocimum sanctum* is a great option for the environmentally friendly manufacture of silver nanoparticles since it contains beneficial components such flavonoids, rosmarinic acid, and eugenol [16]. These phytochemicals can function as capping agents, stabilizing the nanoparticles and avoiding agglomeration, and reducing agents [18 & 19], transforming silver ions into silver nanoparticles. Although the effectiveness of several plant extracts in the production of nanoparticles has been shown in earlier research [20], *Ocimum sanctum*’s unique potential is still not fully understood. Investigating this potential not only contributes to the field of green nanotechnology but also enhances the value of *Ocimum sanctum*, which is already well-regarded in traditional medicine.

Using *Ocimum sanctum* leaf extract as a bioreductant, this study attempts to synthesize silver nanoparticles and analyze the final products using several analytical methods. The successful synthesis and stability of AgNPs was verified using Fourier-transform Infrared spectroscopy (FTIR), Scanning Electron Microscopy (SEM), X-ray diffraction (XRD), and UV-vis spectroscopy. By identifying the surface plasmon resonance, a distinctive optical characteristic of metal nanoparticles, AgNP production and stability are tracked using UV-vis spectroscopy. The crystalline structure and average size of the nanoparticles are revealed by XRD analysis [21]. FTIR spectroscopy assists in identifying the functional groups involved in the reduction and stabilization process [12, 17 & 22], while SEM provides comprehensive imaging to analyze the shape and size distribution of the nanoparticles [23].

Additionally, the antimicrobial efficacy of the biosynthesized nanoparticles was evaluated to establish their potential application in antimicrobial treatments. The antibacterial activity was assessed against *Escherichia coli*, measuring the inhibition zones to determine the effectiveness of AgNPs. Understanding the dose-dependent response of microbial inhibition by silver nanoparticles is crucial for optimizing their application in medical and environmental settings [28]. The successful synthesis of silver nanoparticles using *Ocimum sanctum* not only provides a green alternative to traditional methods but also underscores the potential of integrating traditional medicinal plants with modern nanotechnology [24 & 25]. By advancing our understanding of green synthesis techniques, this research supports the broader goal of developing sustainable technologies that address global challenges in health and the environment [26 & 27].

## Materials and Methods

### Materials

Analytical grade silver nitrate (AgNO_4_) was purchased from a licensed vendor. We picked fresh tulsi (*Ocimum sanctum*) leaves from a nearby garden. Analytical grade ethanol was used to wash the plant leaves, and distilled water was used for all preparations and dilutions. Before being utilized in the experiment, every piece of glassware was carefully cleaned and rinsed with distilled water.

### Preparation of *Ocimum sanctum* Extract

After gathering fresh *Ocimum sanctum* leaves, they were carefully cleaned with distilled water to get rid of any dust or other contaminants. After rinsing the leaves in ethanol to guarantee that all microbiological pollutants were eliminated, they were set to air dry at ambient temperature. A 250 mL Erlenmeyer flask with 100 mL of distilled water was filled with about 20 grams of finely chopped dry leaves. The bioactive components in this mixture were extracted by heating it to 60°C for 30 minutes. Whatman No. 1 filter paper was used to filter the extract once it had cooled to room temperature in order to get rid of any solid residues. For later usage, the filtrate was gathered and kept in storage at 4°C.

### Synthesis of Silver Nanoparticles

0.017 grams of silver nitrate (AgNO_4_) were dissolved in 100 milliliters of distilled water to create a 0.001 M solution of AgNO_4_. In order to create silver nanoparticles, 90 mL of 0.001 M AgNO_3_ solution and 10 mL of *Ocimum sanctum* leaf extract were combined in a 250 mL Erlenmeyer flask. At room temperature, the reaction mixture was continually swirled. The solution’s hue changed from pale yellow to brown, signifying the creation of silver nanoparticles. In order to guarantee total reduction of Ag+ ions, the reaction was allowed to continue for a whole day.

### Characterization of Silver Nanoparticles

By analyzing the UV-vis spectra of the reaction media at various time intervals (0, 1, 2, 4, 8, 12, and 24 hours), the reduction of pure Ag ions was observed. Using a UV-vis spectrophotometer, the spectra was captured in the 200–800 nm region.

Using X-ray diffraction (XRD), the crystalline structure of the produced silver nanoparticles was ascertained. Using a diffractometer, the dried powder of silver nanoparticles was scanned in the 2θ range of 20-80°. The Debye–Scherrer equation was utilized to estimate the average size of the nanoparticles as follows:

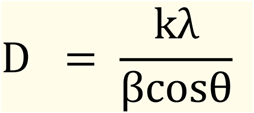

*Where D = Thickness of the nanocrystal*,

*k = Constant (0*.*94)*,

*λ = Wavelength of X-rays*,

*β = Width at half maxima of (111) reflection at Bragg’s angle 2θ*,

*θ = Bragg angle [32]*.

Atomic Absorption Spectroscopy (AAS) analysis of the concentration of silver ions revealed the transformation of Ag ions into Ag nanoparticles. After adding leaf extract, the concentration of Ag ions in the reaction solution was now observed at various intervals. The silver nanoparticles’ size and shape were examined using Scanning Electron Microscopy (SEM). An infrared lamp was used to dry a drop of the nanoparticle suspension that had been applied to a copper grid covered with carbon. After that, an electron microscope was used to image the samples. To determine which functional groups in the *Ocimum sanctum* extract were responsible for the reduction and stability of silver nanoparticles, Fourier-transform infrared (FTIR) spectroscopy analysis was performed. After being combined with potassium bromide (KBr) and pressed into pellets, the dried nanoparticles were examined in the 4000-400 cm^-1^ range with an FTIR spectrophotometer.

### Antimicrobial Activity

The antibacterial activity of the generated silver nanoparticles was assessed against *Escherichia coli*. The bacterial cultures were maintained on nutrient agar slants and subcultured in nutrient broth before to use [38]. Using a sterile cotton swab, a grass culture of each test microorganism was uniformly distributed on the Mueller-Hinton agar plate surface. After inoculating agar plates, sterile paper disks impregnated with different amounts of the suspension of silver nanoparticles were placed on top of them. After the plates were treated for 24 hours at 37°C, the zone of inhibition surrounding each disk was measured to assess the antibacterial activity of the silver nanoparticles.

## Results

### Formation of Silver Nanoparticles

A visual confirmation of the production of silver nanoparticles was obtained when the reaction mixture’s color changed from light yellow to brown [29]. The reduction of silver ions (Ag+) to silver nanoparticles (AgNPs) was shown by this color shift [30].

### UV-vis Spectroscopy of Silver Nanoparticles

UV-vis spectroscopy was used to further establish the AgNPs’ production and stability. A silver nitrate solution’s color changed from translucent to dark yellow upon the addition of *O. sanctum* leaf extract, signifying the creation of silver nanoparticles [29]. The silver nanoparticles’ surface plasmon vibrations are excited, which results in this hue shift [31]. These nanoparticles’ surface plasmon resonance (SPR) produced a peak with a center of 420 nm (figure 1).

**Figure 1:**
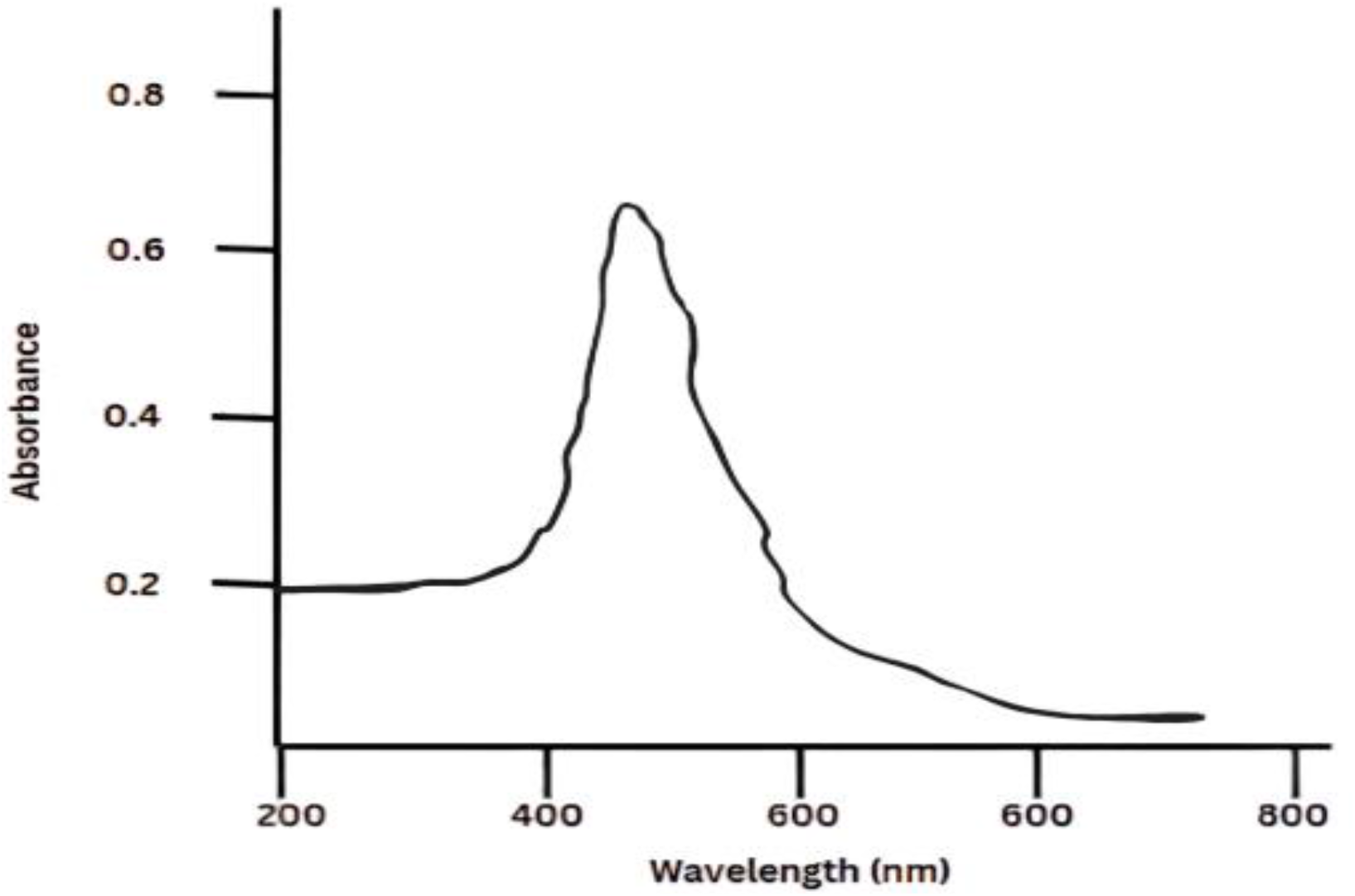
UV-vis Spectrum of Synthesized Silver Nanoparticles from *Ocimum Sanctum*

### X-ray Diffraction (XRD) Analysis

Diffraction peaks corresponding to the silver’s fcc structure were revealed by the XRD study. Figure 2 shows the locations of intense peaks at 37.81, 45.01, 63.91, and 78.0, which correspond to Bragg’s reflections at 111, 200, 220, and 311, respectively. The development of nanoparticles is shown by the broadening of the Bragg peaks. The average particle size was calculated using Scherrer’s formula [37] and full width at half maximum (FWHM) data. It was estimated that the average particle size was about 18 nm.

**Figure 2:**
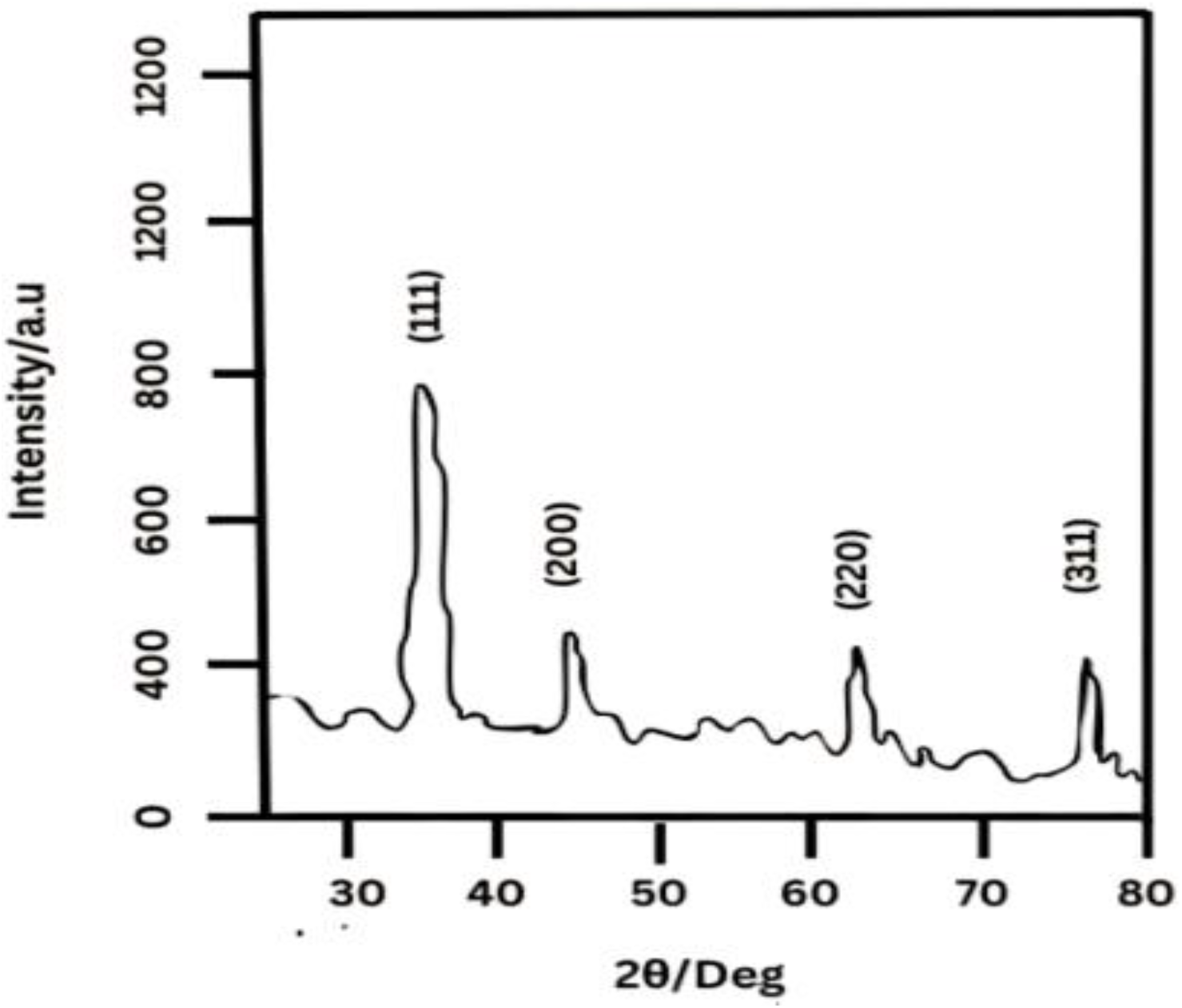
XRD Pattern of Synthesized Silver Nanoparticles from *Ocimum Sanctum*

### Atomic Absorption Spectroscopy (AAS) Analysis

Atomic Absorption Spectroscopy (AAS) was used to detect the concentration of silver ions (Ag ions) in ppm over time during the creation of silver nanoparticles using *Ocimum sanctum* extract. The concentration of Ag ions starts out at 2.73 ppm and quickly drops to 2.06 ppm in the first minute as shown in figure 3. This rapid initial reduction is likely caused by the extract’s active reducing agents. The production of nanoparticles continues as the concentration drops to 1.91 ppm by the second minute. The concentration shows a minor decrease in the decreasing rate at 2.5 minutes, dropping to 1.5 ppm. It drops sharply to 0.55 ppm by the third minute, suggesting a very effective decrease phase. The concentration starts to drop at 3.5 minutes, reaching 0.18 ppm as the reaction is virtually finished. By the fourth minute, it hits 0.023 ppm, suggesting that the reduction of silver ions is practically complete. These findings show how effective *Ocimum sanctum* extract is as a stabilizing and reducing agent, attaining a notable reduction in a brief amount of time, which is beneficial for large-scale production. The quick and significant reduction of silver ions highlights the possibility of employing this green technology to synthesize silver nanoparticles in a scalable and ecologically acceptable manner.

**Figure 3:**
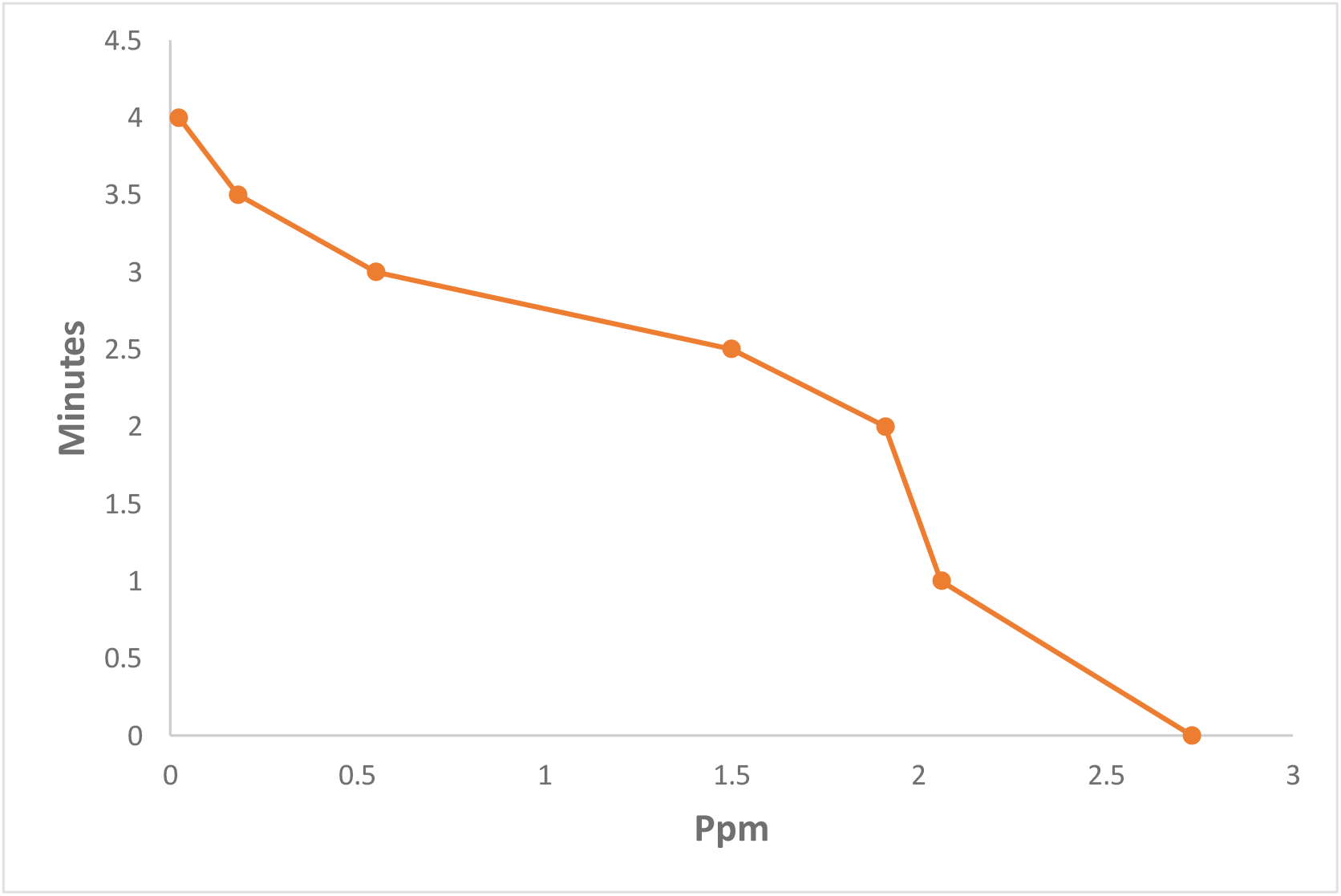
AAS graph of Silver Nitrate Concentration

### Scanning Electron Microscope (SEM) Analysis

Figure 4 presents the scanning electron micrograph of the silver nanoparticles. The image reveals that the particles are spherical, with an average size (mean ± SD) calculated to be 23.82 ± 4.17 nm.

**Figure 4:**
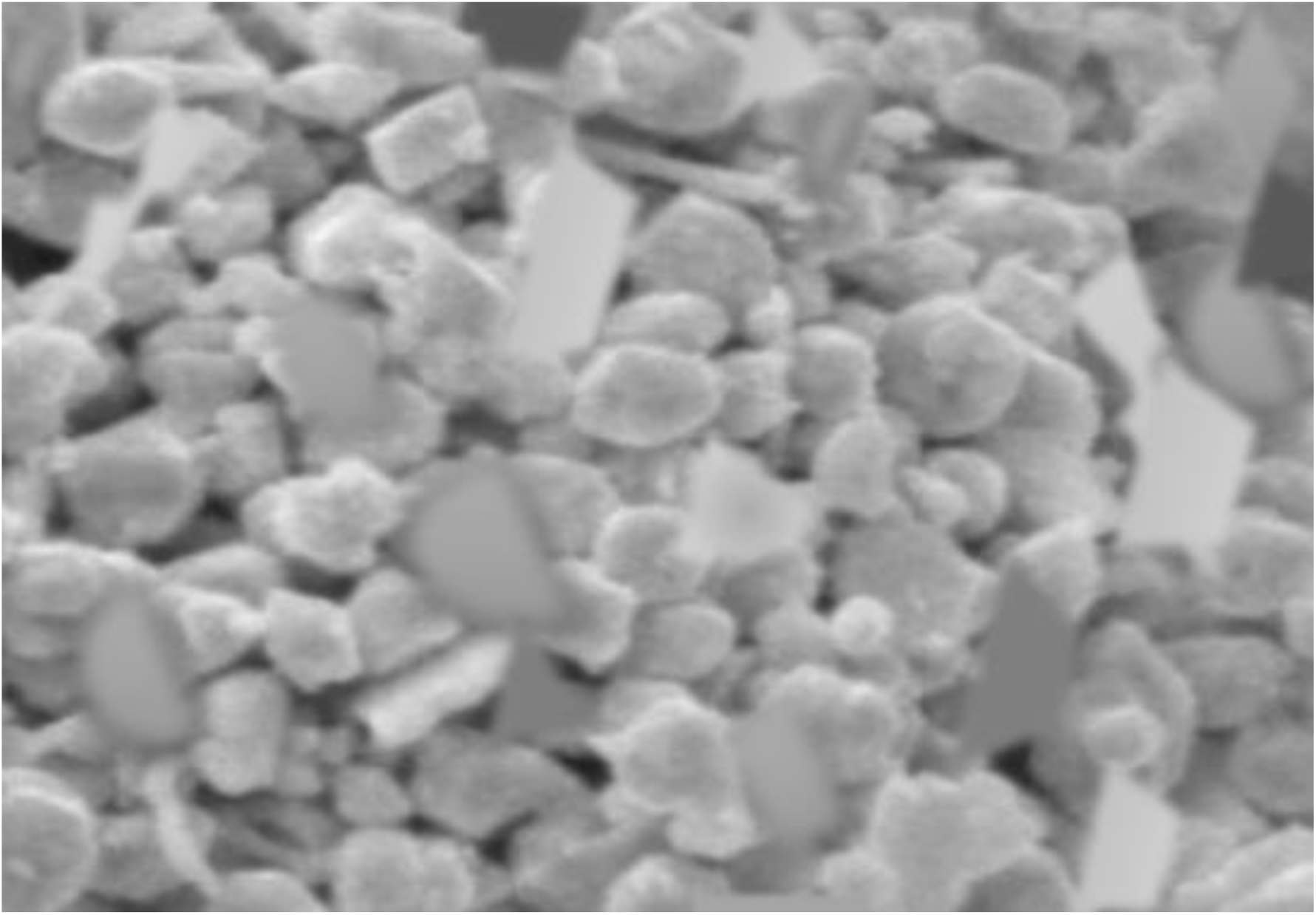
Scanning Electron Micrograph of Silver Nanoparticles

### Fourier Transform Infra-red (FTIR) Analysis

To determine if proteins and silver nanoparticles could interact, FTIR measurements of the biosynthesized particles were performed. The FTIR study’s results revealed distinct absorption peaks at roughly 1635 and 3430 cm^-1^ (Figure 5). According to Jilie and Shaoning (2007), peaks at 3430 cm^-1^ are related to OH stretching in alcohols and phenolic compounds, whereas an absorption peak at 1635 cm^-1^ may be related to the amide I bond of proteins resulting from carbonyl stretch in proteins [34]. It is possible that proteins are interacting with biosynthesized nanoparticles because the absorption peak at 1635 cm^-1^ is similar to that found for native proteins [35] and because the secondary structure of the proteins was unaffected by the reaction with Ag ions or by the binding of Ag nanoparticles with Ag [36]. According to Sathyavathi et al. (2010), these infrared spectroscopy investigations verified that the carbonyl groups in amino acid residues have a potent ability to bind with metal, implying the formation of a layer covering metal nanoparticles and functioning as a capping agent to prevent agglomeration and provide stability to the medium [33]. These findings support the hypothesis that certain proteins function as stabilizing and reducing agents for silver nanoparticles.

**Figure 5:**
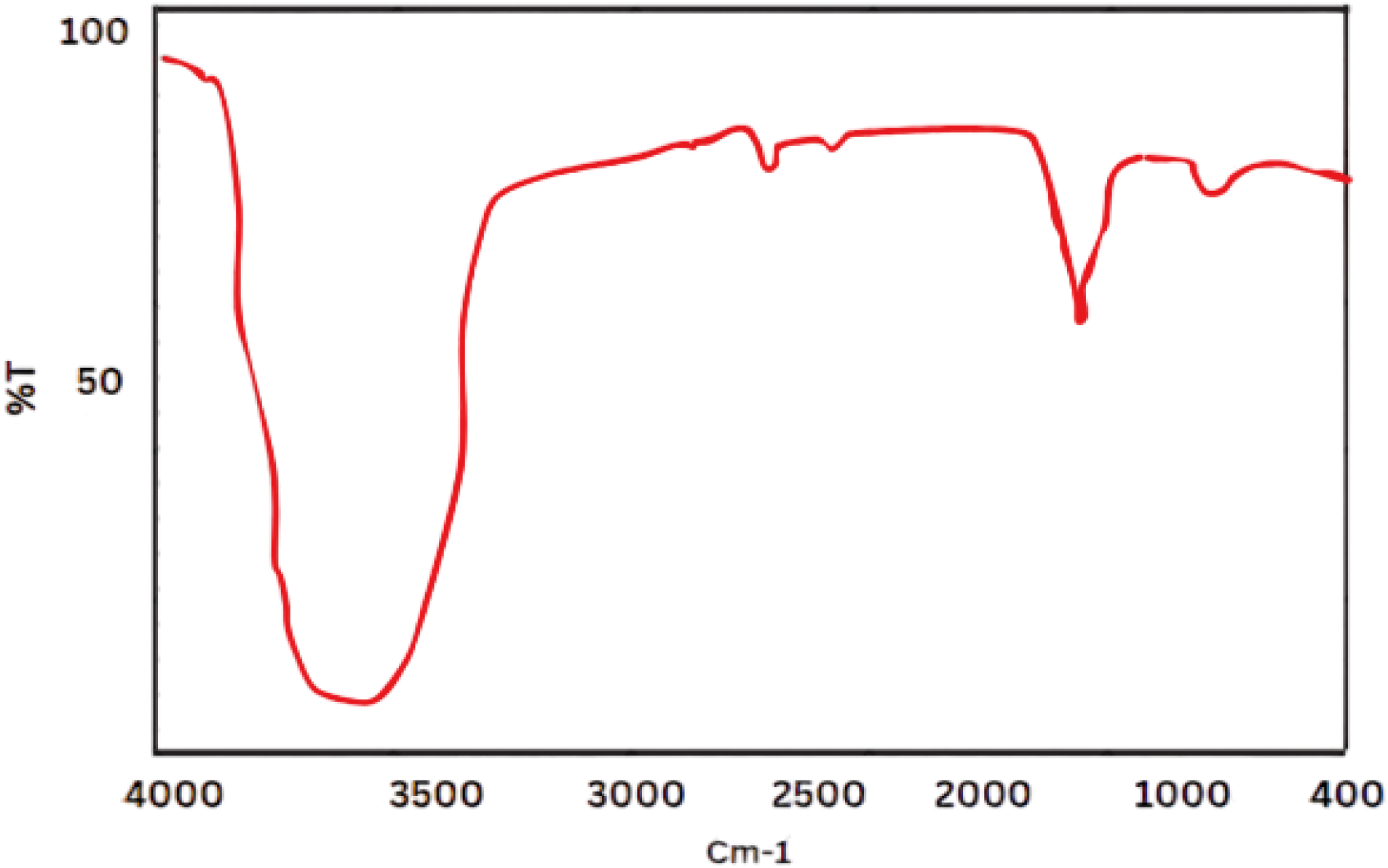
FTIR Pattern of synthesized Silver Nanoparticles

### Antibacterial Activity of Synthesized Silver Nanoparticles

The information in figure6 demonstrates a distinct dose-dependent correlation between the antibacterial activity (measured by the zone of inhibition) and the concentration of biosynthesized silver nanoparticles against *Escherichia coli*. The inhibitory zone measures 6 mm at the lowest dosage of 5 µg/L, suggesting that silver nanoparticles might have antimicrobial effects even in minute amounts. The zone of inhibition grows to 8 mm at a dose of 10 µg/L, indicating increased antimicrobial action. The inhibitory zones measure 10 mm and 12 mm, respectively, at concentrations of 15 µg/L and 20 µg/L, indicating a continuous rise in efficacy.

**Figure 6:**
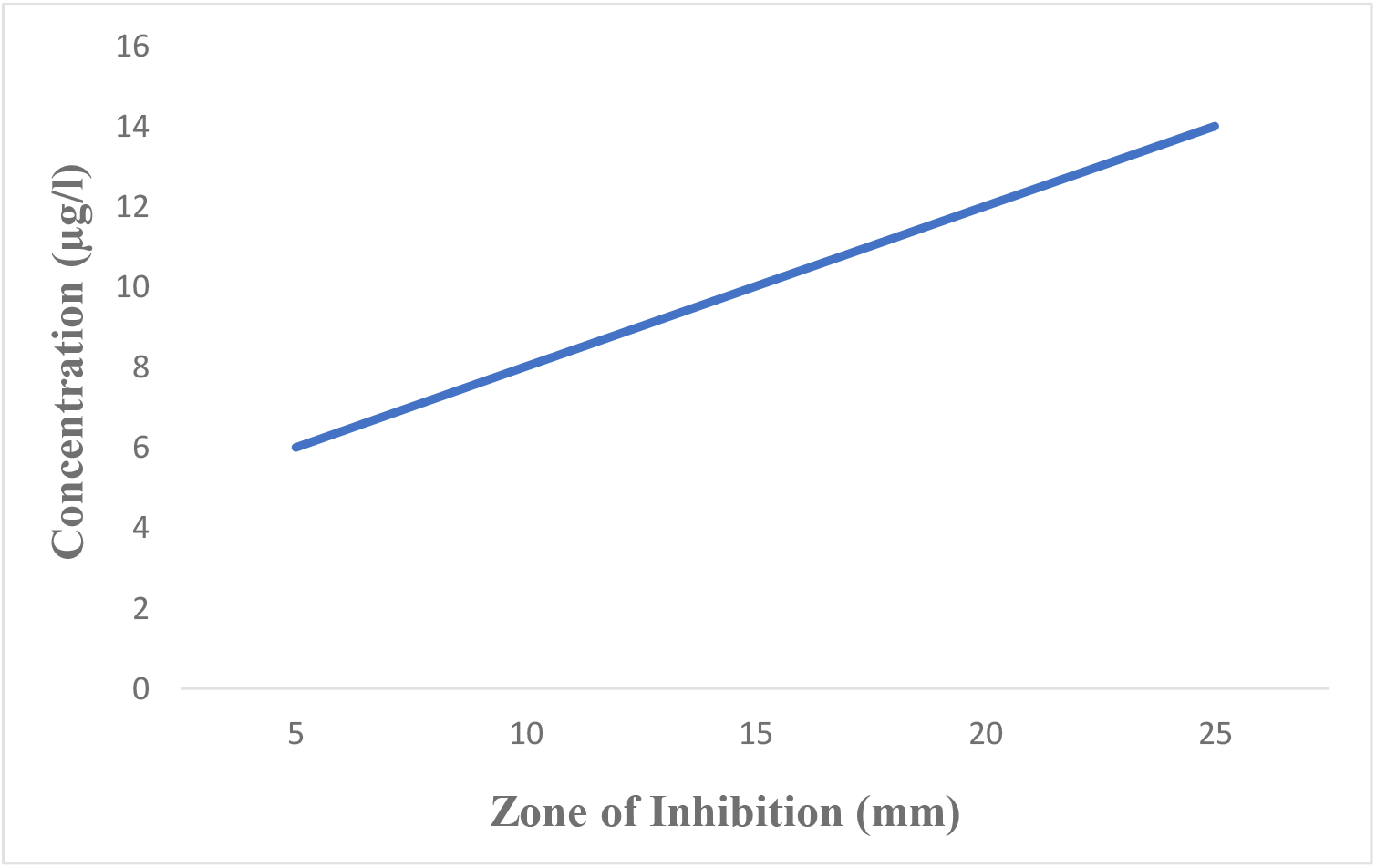
Antibacterial Activity of Silver Nanoparticles against *Escherichia coli*

The inhibition zone reaches 14 mm at the highest dosage of 25 µg/L, indicating a strong antibacterial action. This pattern is consistent with silver nanoparticles’ established mode of action, which includes rupturing microbial cell membranes, producing reactive oxygen species, and interacting with microbial DNA to cause cell death. Higher concentrations of silver nanoparticles show a rise in the inhibitory zone, indicating their potential as powerful antibacterial agents whose activity increases with concentration [24].

## Conclusion

In a nutshell the environmentally sustainable production of silver nanoparticles (AgNPs) by the use of *Ocimum sanctum* (Tulsi) leaf extract has proven to be a feasible substitute for traditional chemical procedures, effectively tackling the dual concerns of environmental sustainability and microbiological resistance. The synthesized AgNPs were confirmed to be stable, have a well-defined crystalline structure, and have a spherical morphology with an average size of about 18– 24 nm by means of UV–vis, X-ray diffraction (XRD), Atomic Absorption Spectroscopy (AAS), Scanning Electron Microscopy (SEM), and Fourier-transform infrared (FTIR) spectroscopy.

Our findings demonstrated the biosynthesized AgNPs’ potential as potent antimicrobial agents by showing a dose-dependent suppression of bacterial growth and strong antimicrobial efficacy. The notable antibacterial potency and the observed fast reduction of silver ions demonstrate the effectiveness of O. sanctum extract as a capping and reducing agent during the nanoparticle manufacturing process. The results of this work provide a viable and sustainable way for producing nanoparticles, and they add to the increasing body of evidence supporting green synthesis techniques. The produced AgNPs’ strong antibacterial qualities point to possible biomedical uses, especially in the creation of fresh antimicrobial therapies and coatings to battle resistant microbial strains.

Future studies should examine these AgNPs’ possible cytotoxicity and biocompatibility in clinical settings, as well as the molecular components of their antibacterial effect. In addition, for industrial applications, it will be essential to scale up the synthesis process while adhering to ecological principles. With wide-ranging effects on the environment and public health, the green production of AgNPs utilizing O. sanctum extract is a noteworthy development in the realm of nanotechnology.

## References

[1]. Naganthran A, Verasoundarapandian G, Khalid F, Masarudin M, Zulkharnain A, Nawawi N, et al. Synthesis, Characterization and Biomedical Application of Silver Nanoparticles. Materials. 2022;15. 10.3390/ma15020427

[2]. Choi C, Schlenker E, Ha H, Cheong J, Hwang B. Versatile Applications of Silver Nanowire-Based Electrodes and Their Impacts. Micromachines. 2023;14. 10.3390/mi14030562

[3]. Ivanišević I. The Role of Silver Nanoparticles in Electrochemical Sensors for Aquatic Environmental Analysis. Sensors (Basel). 2023;23. 10.3390/s23073692

[4]. Sousa M, Guimarães M, Tadielo L, Nascimento B, Rosa D, Oliveira H, et al. Sanitizing activity of silver nanoparticles synthesized with natural products on dairy industry surfaces. Ciência Rural. 2024. 10.1590/0103-8478cr20230027

[5]. Cerqueira I, Mieli M, Pereira A, Corbi P, Resende F. Evaluation of the Cytotoxicity and Genotoxicological Safety Profile of Bioactive Silver(I) Complexes with Aminoadamantane Ligands. Química Nova. 2024. 10.21577/0100-4042.20240002

[6]. Es-Saheb M, Fouad Y, Ibrahim K. Characterization of electrospun pluronic F68/polyvinyl alcohol nanofibers composites containing titanium dioxide, silver, and cobalt nanoparticles for biomedical applications. Materials Express. 2024. 10.1166/mex.2024.2609

[7]. Baláž M, Bedlovičová Z, Daneu N, Siksa P, Sokoli L, Tkáčiková L, et al. Mechanochemistry as an Alternative Method of Green Synthesis of Silver Nanoparticles with Antibacterial Activity: A Comparative Study. Nanomaterials. 2021;11. 10.3390/nano11051139

[8]. Kumar P, Dixit J, Singh A, Rajput V, Verma P, Tiwari K, et al. Efficient Catalytic Degradation of Selected Toxic Dyes by Green Biosynthesized Silver Nanoparticles Using Aqueous Leaf Extract of Cestrum nocturnum L. Nanomaterials. 2022;12. 10.3390/nano12213851

[9]. Gul A, Shaheen A, Ahmad I, Khattak B, Ahmad M, Ullah R, et al. Green Synthesis, Characterization, Enzyme Inhibition, Antimicrobial Potential, and Cytotoxic Activity of Plant Mediated Silver Nanoparticle Using Ricinus communis Leaf and Root Extracts. Biomolecules. 2021;11. 10.3390/biom11020206

[10]. Balčiūnaitienė A, Liaudanskas M, Puzerytė V, Viškelis J, Janulis V, Viškelis P, et al. Eucalyptus globulus and Salvia officinalis Extracts Mediated Green Synthesis of Silver Nanoparticles and Their Application as an Antioxidant and Antimicrobial Agent. Plants. 2022;11. 10.3390/plants11081085

[11]. Mustapha S, Habib MU, Ma’aruf AM, Mustapha A, Shehu H, et al. Sustainable Technique for Neem (Azadirachta Indica) Seed Oil Extraction: Optimization and Characterization. African Journal of Environment and Natural Science Research. 2024;7(2):218–228. 10.52589/AJENSR-5H1FVLHR

[12]. Sulaiman M, Inuwa DI, Aliyu M. Synthesis of silica nanoparticles from sugarcane waste: Precipitation-based size control and characterization. FUDMA Journal of Sciences. 2024;8(3):222–227. 10.33003/fjs-2024-0803-2425

[13]. Chaiyana W, Punyoyai C, Sriyab S, Prommaban A, Sirilun S, Maitip J, et al. Anti-inflammatory and antimicrobial activities of fermented Ocimum sanctum Linn. extracts against skin and scalp microorganisms. Chemistry & Biodiversity. 2021;19. 10.1002/cbdv.202100799

[14]. Kondapalli N, Hemalatha R, Uppala S, Yathapu S, Mohammed S, Surekha M, et al. Ocimum sanctum, Zingiber officinale, and Piper nigrum extracts and their effects on gut microbiota modulations (prebiotic potential), basal inflammatory markers and lipid levels: oral supplementation study in healthy rats. Pharmaceutical Biology. 2022;60:437–450. 10.1080/13880209.2022.2033797

[15]. Chaudhary A, Sharma S, Mittal A, Gupta S, Dua A. Phytochemical and antioxidant profiling of Ocimum sanctum. Journal of Food Science and Technology. 2020;57:3852–3863. 10.1007/s13197-020-04417-2

[16]. Munir N, Khilji S, Shabir M, Sajid Z. Exogenous application of ascorbic acid enhances the antimicrobial and antioxidant potential of Ocimum sanctum L. grown under salt stress. Journal of Food Quality. 2021. 10.1155/2021/4977410

[17]. Sulaiman M, Inuwa DI, Aliyu M. Controlled synthesis of silica particles using extraction-precipitation method. International Journal of New Chemistry. 2024. 10.22034/ijnc.2023.2015271.1364

[18]. Ma’aruf MA, Mustapha S, Giriraj T, Muhammad NS, Habib MU, et al. Sustainable synthesis strategies: Biofabrication’s impact on metal and metal oxide nanoparticles. African Journal of Environment and Natural Science Research. 2024;7(2):229–252. 10.52589/ajensrjtfpyhuk

[19]. Inuwa DI, Aliyu M, Sulaiman M. Evaluation of heavy metals in soil from automobile mechanic village Dutse, Jigawa State, Nigeria. Dutse Journal of Pure and Applied Sciences. 2023;9(2):153–164. 10.4314/dujopas.v9i2a.15

[20]. Ramamurthy J, Jayakumar N. Anti-inflammatory, anti-oxidant effect and cytotoxicity of Ocimum sanctum intra oral gel for combating periodontal diseases. Bioinformation. 2020; 16:1026–1032. 10.6026/973206300161026

[21]. Nath D, Singh F, Das R. X-ray diffraction analysis by Williamson-Hall, Halder-Wagner and size-strain plot methods of CdSe nanoparticles-a comparative study. Materials Chemistry and Physics. 2020;239:122021. 10.1016/J.MATCHEMPHYS.2019.122021

[22]. Miditana S, Tirukkovalluri S, Raju I, Alim S. Photocatalytic degradation of Orange-II by surfactant assisted Mn/Mg co-doped TiO2 nanoparticles under visible light irradiation. Current Chemistry Letters. 2024. 10.5267/j.ccl.2023.6.003

[23]. Melese A, Wubet W, Hussen A, Mulate K, Hailekiros A. A review on biogenic synthesized zinc oxide nanoparticles: synthesis, characterization, and its applications. Reviews in Inorganic Chemistry. 2024. 10.1515/revic-2023-0022

[24]. Dakal T, Kumar A, Majumdar R, Yadav V. Mechanistic Basis of Antimicrobial Actions of Silver Nanoparticles. Frontiers in Microbiology. 2016;7. 10.3389/fmicb.2016.01831

[25]. Muhammad MA, Sulaiman M, Usman HM, Panda SL, Idris IM. Water quality assessment and health implications: A study of Kano metropolis, Nigeria. Journal of Science and Technology. 2024;9(6):33–52. 10.46243/jst.2024.v9.i6.pp33-52

[26]. Muhammad MA, Sulaiman M, Usman HM, Panda SL, Idris IM. Water quality assessment and health implications: A study of Kano metropolis, Nigeria. Journal of Science and Technology. 2024;9(6):33–52. 10.21203/rs.3.rs-4600212/v1

[27]. Usman HM, Sulaiman M, Yahaya MS, Saminu S. Stabilizing environmental conditions for improved biogas generation: A comparative analysis of above-ground and underground plastic digester in fed-batch systems. Iranica Journal of Energy & Environment. 2024.

[28]. Sousa M, Guimarães M, Tadielo L, Nascimento B, Rosa D, Oliveira H, et al. Sanitizing activity of silver nanoparticles synthesized with natural products on dairy industry surfaces. Ciência Rural. 2024. 10.1590/0103-8478cr20230027

[29]. Traoré N, Spruck C, Uihlein A, Pflug L, Peukert W. Targeted color design of silver-gold alloy nanoparticles. Nanoscale Advances. 2024. 10.1039/d3na00856h

[30]. Kumari S, Singh V, Singh D. Nanoparticle synthesis advancements and their application in wastewater treatment: A comprehensive review. Current Chemistry Letters. 2024. 10.5267/j.ccl.2023.9.002

[31]. Mulvaney P. Surface plasmon spectroscopy of nanosized metal particles. Langmuir. 1996; 12:788–800.

[32]. H. M. Usman, M. S. Yahaya, S. Saminu, M. Muhammad, S. Ibrahim, B. I. Sani, M. Sulaiman. Design Modification and Performance Evaluation of Solar PV System at Autocad Laboratory. Leveraging Artificial Intelligence to Achieve United Nations Sustainable Development Goals. Jun. 2 - 6, 2024.

[33]. Sathyavathi R, Krishna MB, Rao SV, Saritha R, Rao DN. Biosynthesis of silver nanoparticles using Coriandrum sativum leaf extract and their application in nonlinear optics. Advanced science letters. 2010 Jun 1;3(2):138–43. 10.1166/asl.2010.1099

[34]. Kong J, Yu S. Fourier transform infrared spectroscopic analysis of protein secondary structures. Acta biochimica et biophysica Sinica. 2007 Aug;39(8):549–59. 10.1111/j.1745-7270.2007.00320.x

[35]. Macdonald ID, Smith WE. Orientation of cytochrome c adsorbed on a citrate-reduced silver colloid surface. Langmuir. 1996 Feb 7;12(3):706–13. 10.1021/la950256w

[36]. Fayaz AM, Balaji K, Girilal M, Yadav R, Kalaichelvan PT, Venketesan R (2010) Biogenic synthesis of silver nanoparticles and their synergistic effect with antibiotics: a study against gram-positive and gram-negative bacteria. Nanomed Nanotechnol Biol Med 6:103–109. 10.1016/j.nano.2009.04s.006

[37]. Klug H.P., Alexander L.E. X-ray Diffraction Procedures for Polycrystalline and Amorphous Materials. Journal of Chemical Education. 1955; 32 (-) :228–235. https://sid.ir/paper/628669/en

[39]. Mustapha Sulaiman, Amina Lawan Abubakar, Ma’aruf Abdulmumin Muhammad et al. Medicinal Assessment of Kenaf (Hibiscus cannabinus L.): Antibacterial, Phytochemical, and Nutritional Profiling, 22 July 2024, PREPRINT (Version 1) available at Research Square 10.21203/rs.3.rs-4783541/v1

